# massNet: integrated processing and classification of spatially resolved mass spectrometry data using deep learning for rapid tumor delineation

**DOI:** 10.1101/2021.05.06.442938

**Authors:** Walid M. Abdelmoula, Sylwia Stopka, Elizabeth C. Randall, Michael Regan, Jeffrey N. Agar, Jann N. Sarkaria, William M. Wells, Tina Kapur, Nathalie Y.R. Agar

## Abstract

**Motivation:** Mass spectrometry imaging (MSI) provides rich biochemical information in a label-free manner and therefore holds promise to substantially impact current practice in disease diagnosis. However, the complex nature of MSI data poses computational challenges in its analysis. The complexity of the data arises from its large size, high dimensionality, and spectral non-linearity. Preprocessing, including peak picking, has been used to reduce raw data complexity, however peak picking is sensitive to parameter selection that, perhaps prematurely, shapes the downstream analysis for tissue classification and ensuing biological interpretation.

**Results:** We propose a deep learning model, massNet, that provides the desired qualities of scalability, non-linearity, and speed in MSI data analysis. This deep learning model was used, without prior preprocessing and peak picking, to classify MSI data from a mouse brain harboring a patient-derived tumor. The massNet architecture established automatically learning of predictive features, and automated methods were incorporated to identify peaks with potential for tumor delineation. The model’s performance was assessed using cross-validation, and the results demonstrate higher accuracy and a 174-fold gain in speed compared to the established classical machine learning method, support vector machine.

**Availability and Implementation:** The code is publicly available on GitHub.

## Introduction

Mass spectrometry imaging (MSI) is a rapidly growing technology that can provide spatial mapping of a wide range of biomolecular classes (such as proteins, metabolites, and lipids) simultaneously and directly from a tissue section in a label-free manner (McDonnell and Heeren, 2007). These make MSI a promising discovery tool with the potential to impact the accuracy and speed of cancer diagnosis and complement the current practice of anatomic pathology (Norris and Caprioli, 2013; Dewez *et al*., 2020). In support providing information that can complement anatomic pathology, MSI can detect molecular alterations in diseased tissues prior to the manifestation of observable morphological changes (Addie *et al*., 2015; Abdelmoula *et al*., 2016; Randall *et al*., 2020). Recent developments have also enabled automated multi-modal integration between MSI and histology (Abdelmoula *et al*., 2014; Patterson *et al*., 2018; Race *et al*., 2021). Such multi-modal integration can harness complementary molecular and anatomical information from biological systems, enabling, for example, a better understanding of molecular mechanisms and pathology of various diseases (Chaurand *et al*., 2004; Veselkov *et al*., 2014), more informed drug metabolite distribution (Castellino *et al*., 2011; Randall *et al*., 2018), and more sensitive surgical guidance (Santagata *et al*., 2014; Eberlin *et al*., 2014).

Matrix-assisted laser desorption ionization (MALDI) is a promising sample introduction technique for the development of diagnostic applications based on mass spectrometry imaging (Calligaris *et al*., 2015; Drake *et al*., 2017; Basu *et al*., 2019; Huizing *et al*., 2019). Acquisition of spectral data from different molecular classes depends largely on the tissue preparation and matrix selection (Carreira *et al*., 2015). MSI data acquisition is done through laser scanning of a tissue surface for desorption and ionization of molecules using a predefined spatial resolution grid, which defines pixel dimensions. The mass-to-charge ratio (*m/z*) and relative intensity of the released ions are detected by a mass analyzer. Each pixel then provides a mass spectrum that would be considered a high dimensional data point for machine learning purposes. The quality and complexity of mass spectral data depend upon the mass analyzer. Fourier-transform ion cyclotron resonance mass spectrometry imaging (FT-ICR MSI) offers the highest mass resolving power (Bowman *et al*., 2020). Such ultra-high mass resolution technology significantly improves the mass identification accuracy, but at the cost of concomitant increases in data complexity and volume (L A McDonnell *et al*., 2010).

The complexity of raw MSI data poses challenges for classical machine learning approaches, where prone to the curse of dimensionality-related issues and slow processing (Alexandrov, 2020). These complexities can be described in terms of massive dimensionality (e.g. 10^5^ – 10^7^ *m/z* values) and a large number of spectra—that can exceed one million spectra per image, depending on both spectral and spatial resolutions—which can reduce clustering and classification quality (Van Der Maaten *et al*., 2009). MSI data preprocessing such as peak picking has been a fundamental step that, to avoid the aforementioned limitations, preceded the analysis routines of classical machine learning (L A McDonnell *et al*., 2010). This fundamental preprocessing step aims to significantly reduce the original data dimensionality through identification of as many relevant peaks as possible while minimizing the noise. However, the available implementations of currently established peak picking approaches are highly sensitive to the selection of parameters that require expert optimization (*e*.*g*. peak shape, signal-to-noise ratio, Full-Width-Half-Maximum, peak frequency, baseline subtraction and spectral smoothing) (Donnelly *et al*., 2019; Murta *et al*., 2021). In addition, the computational performance of peak picking approaches implemented in widely used commercial software is typically quadratic (Alexandrov, 2012), resulting in slow analysis that can take several hours or even a few days depending on mass resolving power and the number of mass spectra processed. A faster alternative but less sensitive approach is to base the peak picking analysis on the mean spectrum (Alexandrov, 2012). McDonnell et al. showed that peak picking on the base peak spectrum, which displays the maximum intensity of each *m/z* value across the dataset, could improve results compared to the analysis of the mean spectrum (Liam A. McDonnell *et al*., 2010), although this analysis is still depended upon expert optimization of peak picking parameters. Spectral preprocessing and the effect of parameters selection do not only impact the downstream analysis (e.g. dimensionality reduction, clustering and classification) but can also affect the overall biological interpretation (Seddiki *et al*., 2020; Murta *et al*., 2021).

Deep learning methods have revolutionized many application areas of biomedical imaging (Ronneberger *et al*., 2015; Esteva *et al*., 2017; Hosny *et al*., 2018). Unlike well-established classical approaches that require feature engineering (Calligaris *et al*., 2015; Balluff *et al*., 2015), deep learning can perform automatic learning of predictive features (Lecun *et al*., 2015). Deep learning techniques offer scalability, non-linearity and efficiency that can accommodate the complexity of MSI data (Abdelmoula *et al*., 2020; Thomas *et al*., 2016; Inglese *et al*., 2017). For example, convolutional neural networks (CNN) were successfully applied to preprocessed and peak picked MSI data for tumor classification (Behrmann *et al*., 2018; van Kersbergen *et al*., 2019; Guo *et al*., 2020). Recently, CNN methods revealed promising results when applied on small scale MSI data without prior processing and (Seddiki *et al*., 2020). Despite the promising results of these CNN architectures, the classification process is sensitive to a user defined, hyper parameter at the input layer that is referred to as the receptive field. The receptive field defines a convolutional kernel window in these CNN architectures to identify salient mass spectral patterns that depend on the selected size of the receptive field (Behrmann *et al*., 2018). Fully connected neural networks (FCNN) were applied on MSI data to perform non-linear dimensionality reduction (Thomas *et al*., 2016; Inglese *et al*., 2017; Dexter *et al*., 2020), and we recently applied FCNN-based architecture to capture spatial patterns and learn underlying *m/z* peaks of interest from large scale MSI data while bypassing conventional preprocessing (Abdelmoula *et al*., 2020).

In this work, we extend our recent deep learning development of msiPL (Abdelmoula *et al*., 2020) to enable tissue classification while avoiding potential bias from the user’s parameter optimization in spectral preprocessing. We propose, **massNet**, a scalable deep learning architecture to perform probabilistic pixel-based classification directly from mass spectral data with massive dimensionality (e.g. tens of thousands of *m/z* values) and without prior preprocessing such as peak picking. Unlike classical machine learning methods, massNet is capable of automatically learning predictive features from large scale MSI data. We demonstrate our method on a MALDI 9.4 Tesla FT-ICR MSI dataset from a patient-derived xenograft (PDX) mouse brain tumor model of glioblastoma (GBM). The model stability was evaluated using 5-fold cross validation, and the classification speed and robustness were assessed on a test set using various established machine learning metrics such as receiver operating characteristic curve (ROC), accuracy, recall, precision and F1-score. Moreover, the classification performance of massNet was benchmarked by comparison to support vector machine (SVM) a widely-used classical machine learning method.

## Materials and Methods

### MALDI FT-ICR MSI Data

MALDI FT-ICR MSI data acquisition was performed on tissue sections from four different intracranial GBM PDX models, see Figure 1. This MSI data and acquisition process were previously published in another study from our group (Randall *et al*., 2020). Briefly, 8 GBM tissue sections of 12 µm thickness were prepared and analyzed using a 9.4 Tesla SolariX mass spectrometer (Bruker Daltonics, Billerica, MA) in the positive ion mode with spatial resolution of 100 µm. The MSI data was exported from SCiLS lab 2020a (Bruker, Bremen, Germany) in the standardized format imzML (Race *et al*., 2012) and converted to the HDF5 format (Folk *et al*., 2011) for deep learning analysis.

**Figure1.**
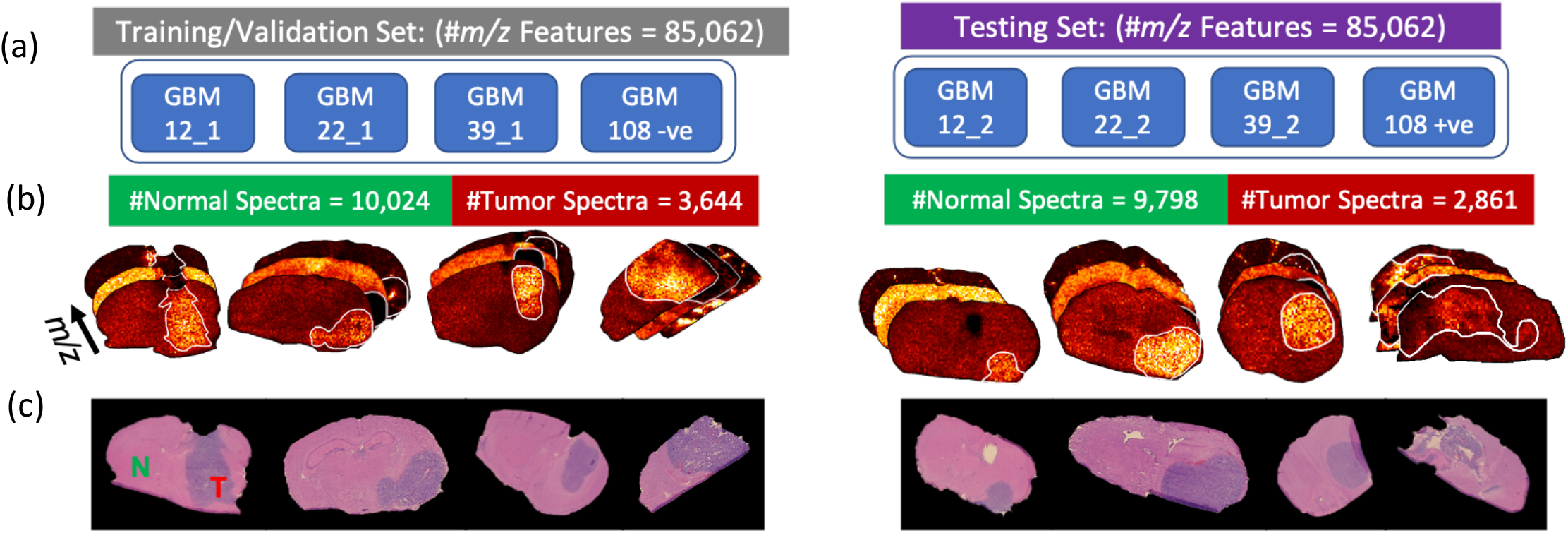
Tissue sections from 4 intracranial GBM PDX models were divided into training/validation and testing sets: (a) schematic distribution of tissue sections from different GBM models, (b) Annotated tumor regions in the MSI datasets which were guided by the H&E annotations (c).

### Microscopy Imaging and Tumor Annotation

Four tissue sections consecutive to those used for the MSI analysis were thaw mounted onto glass slides for hematoxylin and eosin (H&E) staining. The H&E images were acquired using a Zeiss Observer Z.1 microscope equipped with 20x Plan-APOCHROMAT lens and AxioCam MR3 camera. The tumor regions in these H&E images were annotated by an expert pathologist and the annotations were manually transferred to annotate the MSI data as demonstrated in (Randall *et al*., 2020), see Figure 1.

### Deep Learning Architecture of massNet

The hyperspectral image from MSI encompasses a set of high-dimensional data points *X* = {*x*^(1)^, *x*^(2)^, …, *x*^(*N*)^}, where *N* is the total number of spectra (or pixels) and each spectrum is a *d*-dimensional point *x*^(*i*)^ ∈ *R*^(*d*)^ where *d*, is the total number of *m/z* variables. The pixel-wise annotation of the hyperspectral image is represented by a binary vector *Y* = {*y*_1_, *y*_2_, …, *y*_*N*_}, where *y*_*i*_ ∈ [0,1]^*C*^ is the ground truth class label of spectrum *x*^(*i*)^which belongs to one of the total two class labels *C* which are here normal and tumor. Our proposed massNet architecture shown in Figure 2 aims to learn a predictive function ℱ that would establish the relationship between each spectrum *x*^(*i*)^and its associated class label *y*_*i*_ as depicted in equation (1),

**Figure 2.**
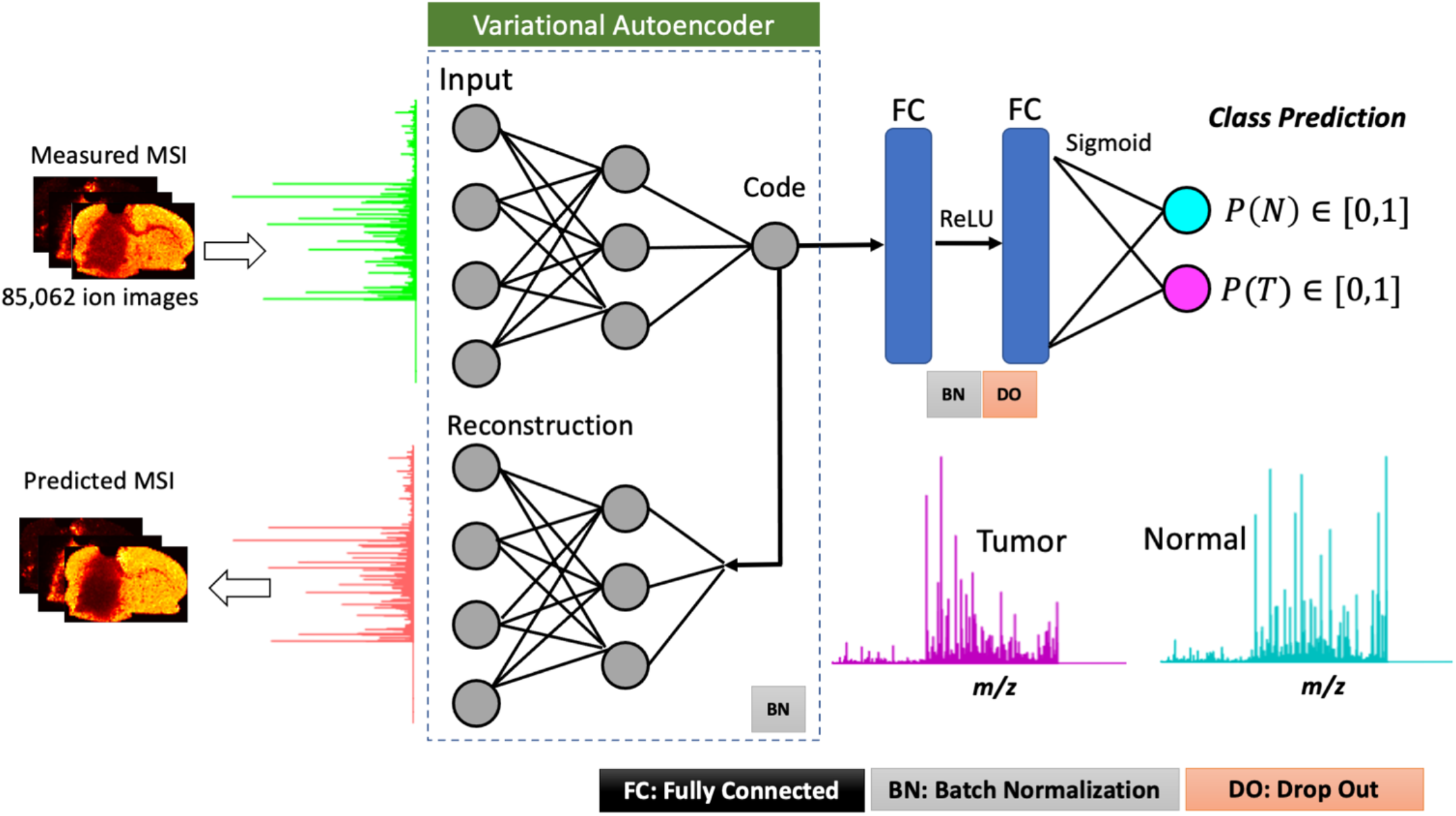
Deep learning-based architecture of massNet for probabilistic two-class classification of large scale MSI data without prior preprocessing and peak picking. The artificial neural network is based on spectral-wise analysis and consists of two modules, namely: variational autoencoder for non-linear manifold learning that is captured at the “Code” layer, and two fully connected layers that take input from the “Code” layer to yield probabilistic predictions at the output layer using the sigmoid activation. massNet is regularized based on batch normalization and drop out to maintain learning stabilization and faster optimization.

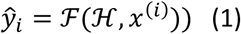

where ŷ_*i*_ ∈ [0,1]^*C*^ is a probabilistic estimate of the predicted class labels *C* that is computed based on the optimized artificial neural network hyper-parameter set ℋ. Class prediction of normal and tumor for a given spectrum *x*^(*i*)^ can then be respectively represented as *P*(*N*) ∈ ŷ_*i*_^*c*=1^ and 1 − *P*(*N*)=*P*(*T*) ∈ ŷ_*i*_^*c*=2^ The massNet architecture has two main modules to optimize the hyper-parameter set ℋ, namely the variational autoencoder (VAE) and probabilistic classification modules. The VAE module aims to learn a non-linear manifold of the original complex MSI data through optimizing two functions: a probabilistic encoder *P*_Ø_(*z*|*x*) and a probabilistic decoder *P* _Ø_ (*z*|*x*)for data reconstruction. The learned non-linear manifold represents a latent variable z ∈*R*^*k*^ ;*k* ≪ *d*, which is of lower dimensions (*k*) and assumed to be sampled from a normal distribution (Coombes *et al*., 2005) as such *z* ∼ *N*(μ_Ø_(*x*), σ_Ø_(*x*)). The VAE parameters Ø and σ are optimized based on maximizing the variational lower bound (ℒ) that is given in equation (2). For more information on VAE we refer readers to Kingma *et al*. (Kingma and Welling, 2013).

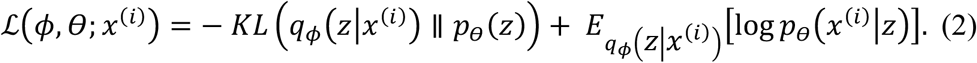

The second module of massNet performs probabilistic classification, which is activated with a sigmoid function 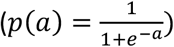, where *a* is an arbitrary variable at the output layer. This classifier module is parametrized by a hyper-parameter ζ and consists of two fully connected hidden layers and an output layer of two classes (normal and tumor), see Figure 2. These two hidden layers are activated based on a rectified linear unit (ReLU) and take their input from the optimized latent variable (*z*) of the VAE. The classification result is based on optimizing a loss function of the binary cross entropy (Ε), shown in equation (3), to maximize the similarity between the real and predicted class labels. Finally, the hyper-parameters ℋ, given in equation (1), consist of hyper-parameters from these two optimized modules ℋ = {Ø, ζ}.

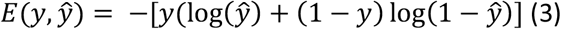

### Data Visualization and Feature Localization

Uniform Manifold Approximation and Projection (UMAP) is a non-linear dimensionality reduction method for data visualization in a single map representation in which it preserves both local and global data structures from higher dimensional space (McInnes *et al*., 2018). UMAP has been found promising in visualizing different types of omics data (Becht *et al*., 2019). The UMAP algorithm was applied on the learned *k* −dimensional latent variable (*z*) to enable data visualization in a two-dimensional space. Of note, applying the UMAP on the latent variable instead of the original hyper-dimensional MSI data is a more efficient way to avoid the curse of dimensionality (Van Der Maaten *et al*., 2009).

The classification predictions were spatially reconstructed to gain insight into the molecular interpretability of the predicted classes. This enabled the predicted class labels to be mapped back into the image space to reveal spatial patterns associated with each of the predicted classes. Pearson correlation was then applied on each of these spatial patterns and the identified peaks from the VAE module using the msiPL method (Abdelmoula *et al*., 2020). The highly co-localized *m/z* peaks are those that achieved the highest Pearson correlation coefficient (*r* ≥ 0.7).

### Model Evaluation

The model’s learning stability was evaluated using 5-fold cross validation on the training set (80% training and 20% validation), and the best model was finally applied and assessed on an independent test set. The classification speed and robustness were assessed on a test set using various established machine learning metrics such as receiver operating characteristic curve (ROC), accuracy, recall, precision and F1-score. Moreover, the classification performance of massNet was benchmarked by comparison to support vector machine (SVM) a widely-used classical machine learning method.

## Results

### Hyperparameters and implementation details of the massNet architecture

The massNet architecture, shown in Figure 2, consists of an input layer (*L*_*in*_), five hidden layers (*h*_1_, *h*_*code*_, *h*_3_, *h*_4_, *h*_5_), and two output layers (*L*_*out1*_,*L*_*out2*_). The input layer (*L*_*in*_) takes its input signal from total-ion-count (TIC) normalized mass spectra. The VAE module has three hidden layers (*h*_1_, *h*_*code*_, *h*_3_) and an output layer (*L*_*out1*_). The lower dimensional latent variable (*z*) is captured at the Code layer (*h*_*code*_), which is then used by the generative model for spectral data reconstruction at the output layer (*L*_*out1*_). The Code layer (*h*_*code*_) also provides an input to the classification module which has two fully connected hidden layers (*h*_4_, *h*_5_) and eventually yields a probabilistic estimate of class prediction at the output layer (*L*_*out2*_). The layer size of each (*L*_*in*_) and (*L*_*out1*_) is equivalent to the number of *m/z* variables, whereas the size of *L*_*out2*_ is equivalent to the number of class labels. The size of hidden layers is a user tunable parameter, and it was empirically set as (*h*_1_ = 512, *h*_*code*_ = 5, *h*_3_ = 512, *h*_4_ = 128, *h*_5_ = 128) in which the VAE parameter settings were adopted from our former msiPL method (Abdelmoula *et al*., 2020). The rectified linear unit (ReLU) was used as an activation function for all layers except for the two output layers (*L*_*out1*_, *L*_*out2*_) which were activated using the sigmoid function. The proposed neural network architecture was regularized using batch normalization and dropout to stabilize the learning process, fasten the convergence, and significantly reduce overfitting (Srivastava *et al*., 2014; Ioffe and Szegedy, 2015). A 20% dropout was added to the two hidden layers of the classification module. The cost function was optimized based on minibatch processing (batch size = 100) using the adaptive stochastic gradient descent method of Adam optimization (with default learning rate = 0.001) and the total number of epochs for VAE and classification modules were respectively 100 and 30. The architecture was implemented in python using the publicly available deep learning platforms of Keras (Chollet, 2017) and tensorflow (Abadi *et al*., 2016), and it was trained on a PC workstation (Windows 10, Intel Xenon 3.3GHz, 64-bit Windows, and NVIDIA TITAN XP Graphics Card).

### UMAP visualization of the VAE latent variable reveals separation between normal and tumor spectra

The deep learning model was optimized on a training set acquired by MALDI FT-ICR MSI from 4 different intracranial GBM PDX models, as demonstrated in the left column of Figure 1. This training set encompasses mass spectra with 85,062 *m/z* variables that were collected from two different classes of normal and tumor tissue types with total number of spectra 10,024 and 3,644 respectively. The VAE model optimization revealed stable convergence distribution in 100 epochs with random shuffling of data batches (Supplementary Figure S1) and a total running time of 55.7 minutes. The optimized model captured the latent variable (*z*) of 5-dimensions at the Code layer (Supplementary Figure S2) which was then used to efficiently reconstruct the original TIC-normalized MSI data with a total mean squared error (MSE) of 5.69 × 10 ^−5^, see Figure 3.a. The UMAP visualization of the latent variable in two-dimensional space revealed separation of mass spectra from normal and tumor tissues as shown in Figure 3.b. The UMAP features were also colored based on different GBM models to reveal inter-tumor heterogeneity and visualizing the learning efficiency in capturing similarities based on the molecular phenotype and assess potential batch effect (Figure 3.b).

**Figure 3.**
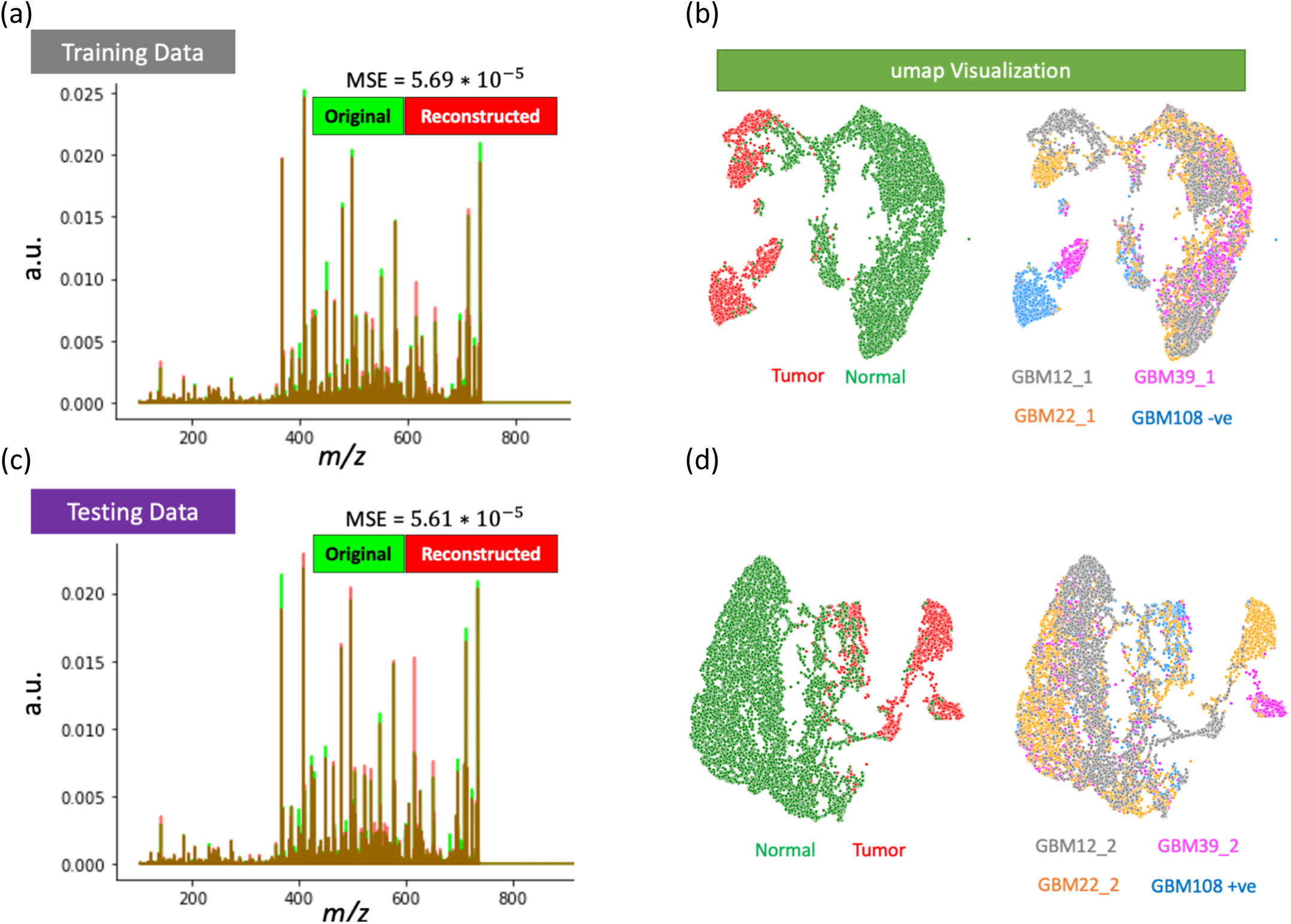
Performance of the VAE module and non-linear data visualization: Overlay of the TIC-normalized average spectrum of original and reconstructed data for both training (a) and testing (b) datasets. UMAP visualization of the 5-dimensional latent variable captured by the VAE model reveals distinction between normal and tumor mass spectra from different GBM models for both training (b) and testing (d) dataset.

### Pre-trained VAE enables optimization of the classification module

The pre-trained and optimized VAE provides input to the classifier module through the latent variable that was captured at the Code layer (*h*_*code*_). The classification module was optimized in 30 epochs which also included random shuffling of data batches and the total running time of 6.43 minutes. To avoid potential overfitting and provide unbiased estimate of out-of-sample error (i.e. on test data), the model was evaluated using 5-fold cross validation. For each of the cross-validation folds, the entire training dataset (shown in Figure 1. a-b) was randomly divided into 80% training and 20% validation. The robustness and evaluation of the different cross-validation models are shown in Supplementary Table S1, in which the best model with maximum accuracy and minimum loss values was eventually selected to be further independently evaluated on the withheld test set. The best model was benchmarked by comparison to SVM where both models showed comparable performance but massNet was faster, see Supplementary Table S2. Of note, the total running time shown in Supplementary Table S2 for optimizing the overall massNet model included the running time for both the VAE and the classification modules.

### Rapid and efficient classification performance on MALDI MSI test set of large dimensions

The optimized massNet model was evaluated on the withheld test set of MALDI FT-ICR MSI data from 4 different intracranial GBM PDX models as demonstrated in the right column of Figure 1. This test set encompasses mass spectra with 85,062 *m/z* variables that were collected from two different classes of normal and tumor tissue types with total number of spectra 9,798 and 2,861 respectively. The optimized VAE module took 61.8 seconds to analyze the entire test set as such the latent variable was captured (Supplementary Figure S3) and the original TIC-normalized MSI test data were reconstructed with an overall MSE of 5.61 × 10^−5^, see Figure 3.c. The UMAP visualization of the 5-dimensional latent variable revealed separation between molecularly distinct phenotypes of normal and tumor tissues, as shown in Figure 3.d.

The optimized massNet took about 29.88 seconds to provide spectral-wise probabilistic prediction of the entire test set. The robustness of this ultra-fast classification (compared to SVM, see Table 1) is supported by ROC analysis (AUC values for normal and tumor classes were 95.01% and 94.94%, respectively) and the confusion matrix which revealed true negative and true positive values of 98% and 83% respectively, as demonstrated in Figure 4. This classification performance was further benchmarked by comparison to an SVM classifier using different machine learning evaluation metrics shown in Table 1. Of note, both the massNet and SVM models were first optimized on the training set and then applied on this test set. The overall performance of massNet was slightly higher than SVM with respective accuracy of 95% and 93.45 %, but more significantly, the massNet was 174 times faster than SVM as shown in Table 1.

**Table 1.** Classification performance on the withheld test MALDI FT-ICR MSI dataset

|  | Class | Spectra # | Precision (%) | Recall (%) | F1-score (%) | Accuracy (%) | Time (min) |
| --- | --- | --- | --- | --- | --- | --- | --- |
| **massNet** | Normal | 9,798 | 95 | 98 | 97 | 95 | 0.5 |
|  | Tumor | 2,861 | 94 | 83 | 88 |  |  |
| **SVM** | Normal | 9,798 | 94 | 98 | 96 | 93.45 | 87.25 |
|  | Tumor | 2,861 | 91 | 79 | 84 |  |  |

**Figure 4.**
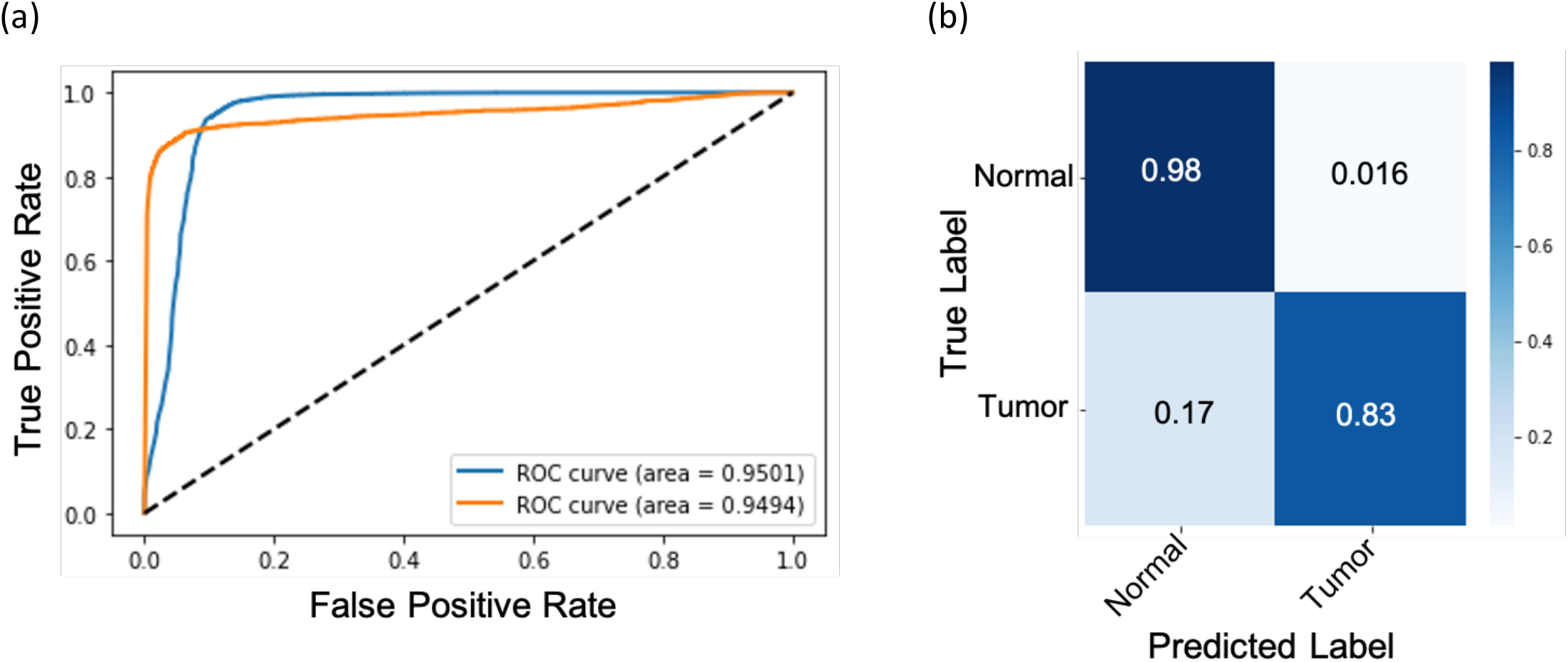
Classification performance on the MALDI FT-ICR MSI test set: (a) Receiver operating characteristic (ROC) curve distribution for both normal (blue) and tumor (orange) classes with an AUC of 95.01% and 94.94%, respectively. (b) Confusion matrix showing the prediction performance compared to the ground truth labels.

### Spatial mapping of predicted scores reveals uncertainty at the tumor margin

The probabilistic scores of the predicted class labels were spatially mapped to enable their visualization in the image space, see Figure 5.a-b. Despite the massNet model being quite accurate in predicting the true class labels in most of the tissue regions, it showed a higher level of uncertainty at the tumor margin (Figure 5.c). This could reflect that the tumor margin has a convoluted molecular signature likely resulting from mixtures of normal and tumor cells compared to either the tumor core or normal tissue types, therefore representing an infiltrative edge. This observation is in accordance with other studies that studied mass spectral signatures from tumor margins (Calligaris *et al*., 2014). Identification of *m/z* peaks that are highly correlated with each of the predicted class labels were determined through the Pearson correlation coefficient. This coefficient was computed by correlating each of the predicted spatial maps with each of the 772 *m/z* values that were identified by msiPL. The top 10 correlated *m/z* values correlated with the tumor region were identified and presented in Table 2, whereas the highest correlated values for normal tissue are shown in Supplementary Table S3. The highest correlated peak with the tumor class was at *m/z* 400.9546 ± 0.01 which is elevated in the tumor region. The automated non-linear image registration method we recently developed (Abdelmoula *et al*., 2014) was applied to non-linearly warp this ion image and fuse it with an image of the Hematoxylin and Eosin (H&E) stained tissue section, Figure 5.d-e. This multi-modal integration can provide a reference system for pathologists to further study and gain more insight about the tissue that can go beyond tissue anatomy (Caprioli, 2019).

**Table 2.** Top 10 correlated m/z values with the tumor tissue

| $m/z$<br>experimental | correlation | Tentative<br>assignment | Molecular<br>formula | Adduct | $m/z$<br>calculated | Database | Error<br>(ppm) |
| --- | --- | --- | --- | --- | --- | --- | --- |
| 400.9546 | 0.8160 | Uridine<br>monophosphate<br>(UMP) | C<sub>9</sub>H<sub>13</sub>N<sub>2</sub>O<sub>9</sub>P | [M+2K-H]<sup>+</sup> | 400.9554553 | HMDB | -2.1 |
| 480.9211 | 0.8104 | Uridine<br>diphosphate<br>(UDP) | C<sub>9</sub>H<sub>14</sub>N<sub>2</sub>O<sub>12</sub>P<sub>2</sub> | [M+2K-H]<sup>+</sup> |  | HMDB | -0.21 |
| 518.8773 | 0.8074 |  |  |  |  |  |  |
| 589.9102 | 0.7909 |  |  |  |  |  |  |
| 545.9591 | 0.7825 | Adenosine<br>triphosphate<br>(ATP) | C<sub>10</sub>H<sub>16</sub>N<sub>5</sub>O<sub>13</sub>P<sub>3</sub> | [M+K]<sup>+</sup> | 545.9589 | Metlin | 0.36 |
| 567.94086 | 0.7746 |  |  |  |  |  |  |
| 464.9471 | 0.7673 |  |  |  |  |  |  |
| 583.91437 | 0.7561 | Adenosine<br>triphosphate<br>(ATP) | C<sub>10</sub>H<sub>16</sub>N<sub>5</sub>O<sub>13</sub>P<sub>3</sub> | [M+2K-H]<sup>+</sup> | 583.9148 | HMDB | -0.74 |
| 703.57416 | 0.7534 | SM (34:1)/PE-Cer (37d/1) | C<sub>39</sub>H<sub>79</sub>N<sub>2</sub>O<sub>6</sub>P | [M+H]<sup>+</sup> | 703.5749 | Metlin | -1.05 |
| 704.5783 | 0.7532 | This is the<br>703.57 C 13<br>isotope |  |  |  |  |  |

**Figure 5.**
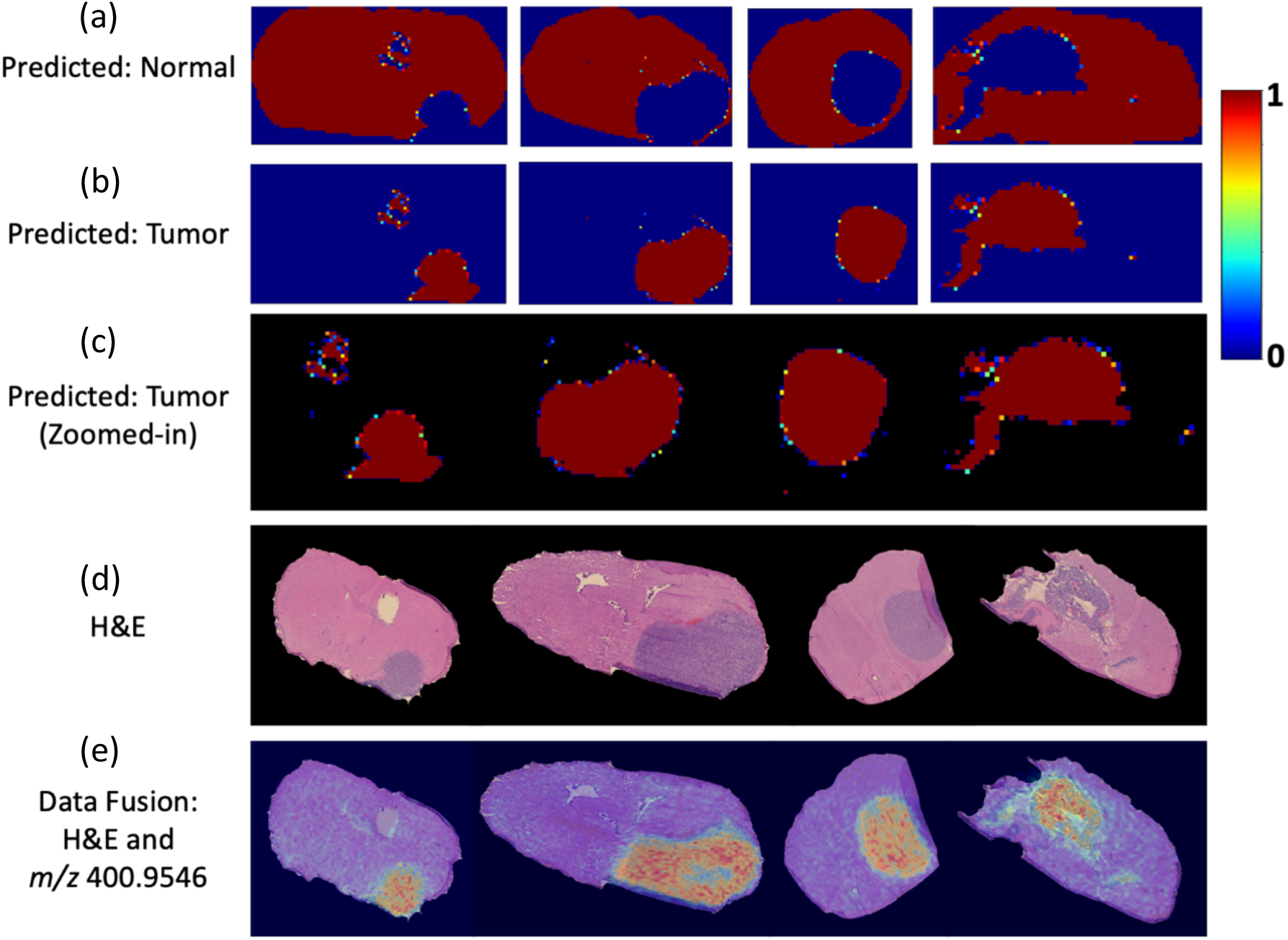
Spatial mapping of the classification predictions and multi-modal integration: (a-b) Spatial distribution of the spectral-wise probabilistic predictions for normal (a) and tumor (b) classes. (c) Close up visualization of the spatially mapped tumor prediction scores reveals a higher level of uncertainty at the interface between normal and tumor (i.e., tumor margins). (d) The H&E microscopy images show tumor regions in different GBM models (columns). (e)Multi-modal integration of the H&E images and ion image at m/z 400.9546 ± 0.01 which is highly correlated and elevated in the tumor region.

## Discussion

We presented massNet, a fully connected deep learning architecture for spectral-wise classification of large scale MSI data without prior preprocessing and peak picking. The massNet architecture consists of two main modules, namely: variational autoencoder (VAE) and classification modules. The VAE module learns a lower dimensional latent variable that was used to reconstruct the original MSI data and assess the learning quality of the VAE module. That optimized latent variable provides an input to the classification modules which was activated at the output layer using a sigmoid function to provide a probabilistic estimate of the predicted class membership. Unlike other architectures for MSI data classification, massNet is based on a fully connected artificial neural network and does not rely on optimizing a receptive field parameter, which is inherently associated with convolutional artificial neural networks and has an impact on the learning process (Behrmann *et al*., 2018; van Kersbergen *et al*., 2019; Guo *et al*., 2020). The optimized model showed high accuracy and ultra-fast performance (less than 1 second) on the full MSI test data that encompassed a total of 12,659 mass spectra with 85,062 *m/z* variables. The trained massNet model achieved higher accuracy and was 174 times faster than a trained SVM when compared for analysis of the same dataset, see Table 1.

While the proposed model showed promising results for binary class classification, the model could be extended in future development to enable multi-class classification. We envision a slight change in the massNet architecture mainly at the output layer (*L*_*out2*_) in which its size will be defined based on the new number of class labels and it must be activated using a different function such as softmax. The ground truth annotations could be defined using microscopy images and then mapped into the MSI space using image registration (Heijs *et al*., 2015). Depending on the application and tissue types, different stainings could be investigated to provide increased structural annotations compared to the most common H&E staining. MSI and microscopy imaging are complementary modalities and annotating the MSI data based solely on annotated histology is an approximation approach which might not be optimal especially in case of intra-tumor heterogeneity (Balluff *et al*., 2015). This is mainly because the MSI data could reveal molecular intra-tumor heterogeneity that is not yet observed in the microscopic image (Randall *et al*., 2020; Jones *et al*., 2012).

The proposed model was applied to the analysis of MALDI FT-ICR MSI data here but the massNet architecture is independent of the mass spectral ionization nature and could be applied to the analysis of MSI data from different platforms with distinct ionization and mass analyzer technologies. The massNet architecture could also be applied to the classification of different tissues and tumor types where the challenging task becomes the establishment of ground truth annotation of the MSI data. For instance, the massNet revealed a higher level of uncertainty around tumor margins in the binary classification task as shown in Figure 5.b-c. This could reflect distinct molecular signatures in the tumor margin area compared to either the tumor core or normal tissue types. Tumor margins could therefore either be treated as an independent class or deconvoluted to extract signal from tumor and normal cells; however, the ground truth annotation is a challenging task that would require more pathological investigation and could be enhanced by integrating with multiplexed immunofluorescence to clearly delineate cell types from the tumor and microenvironment.

## Supplementary Materials

**Figure S1.**
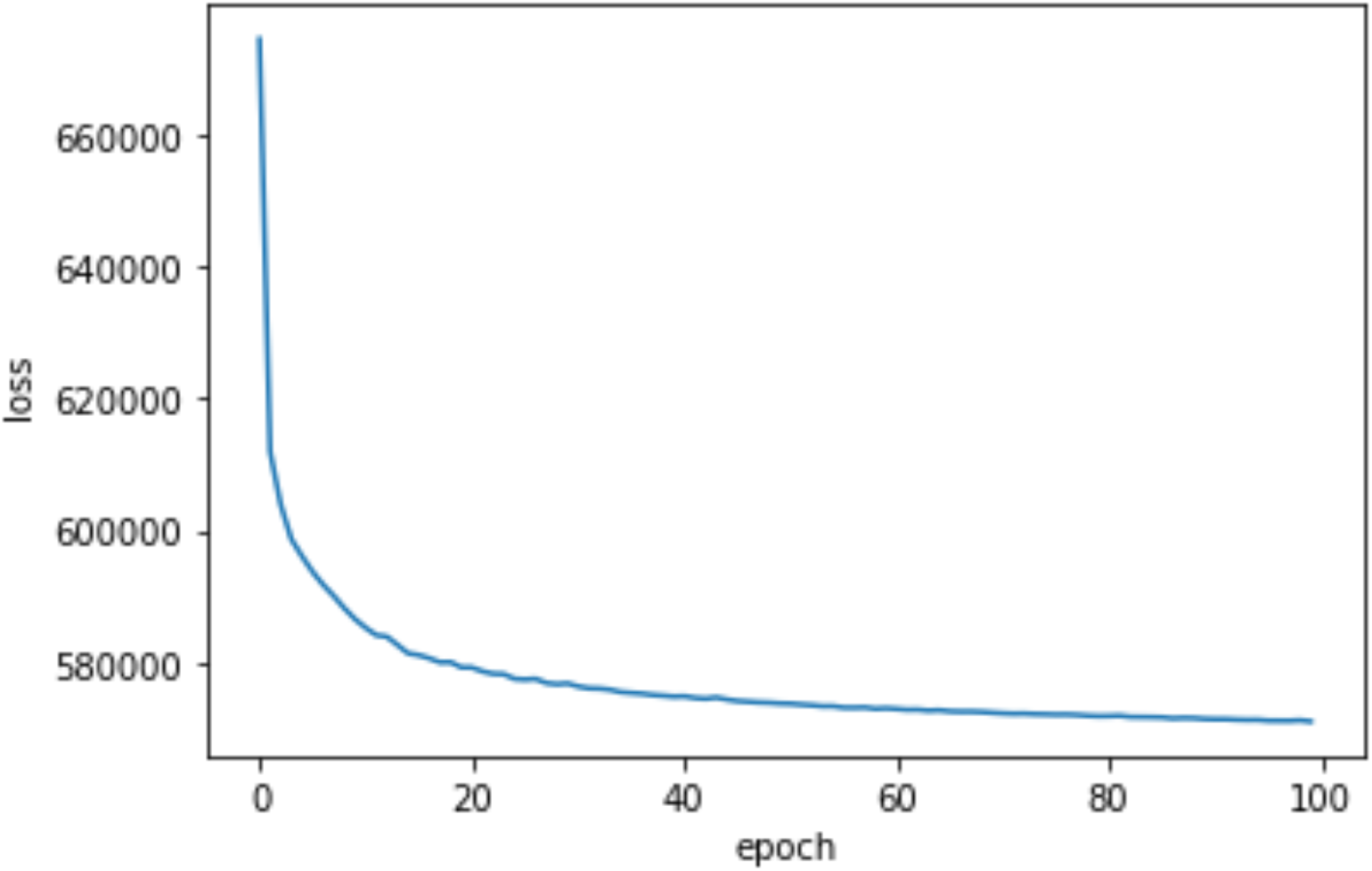
Convergence distribution of the VAE module for 100 epochs (i.e. iterations), and the loss represents the optimization of the VAE’s cost function that consists of a summation of two functions, namely: Kullback-Leibler divergence and binary cross entropy.

**Figure S2.:**
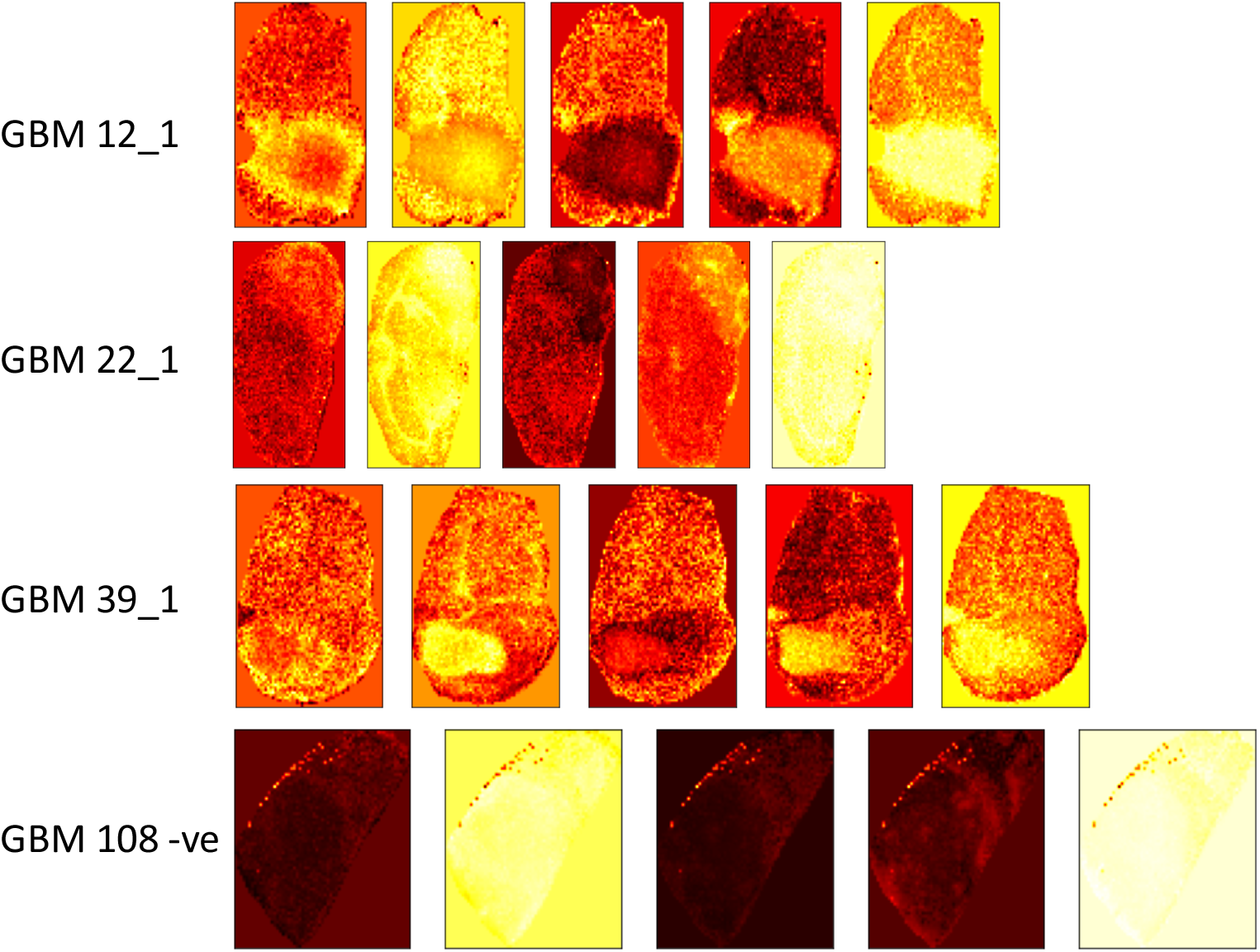
Latent variable of 5-dimensional representation (column) that was captured at the VAE’s code layers for different GBM models (rows) of the MALDI FT-ICR MSI training set. This variable provides a compressed representation of the original MSI data and colormap was arbitrary chosen to reveal structures.

**Figure S3.**
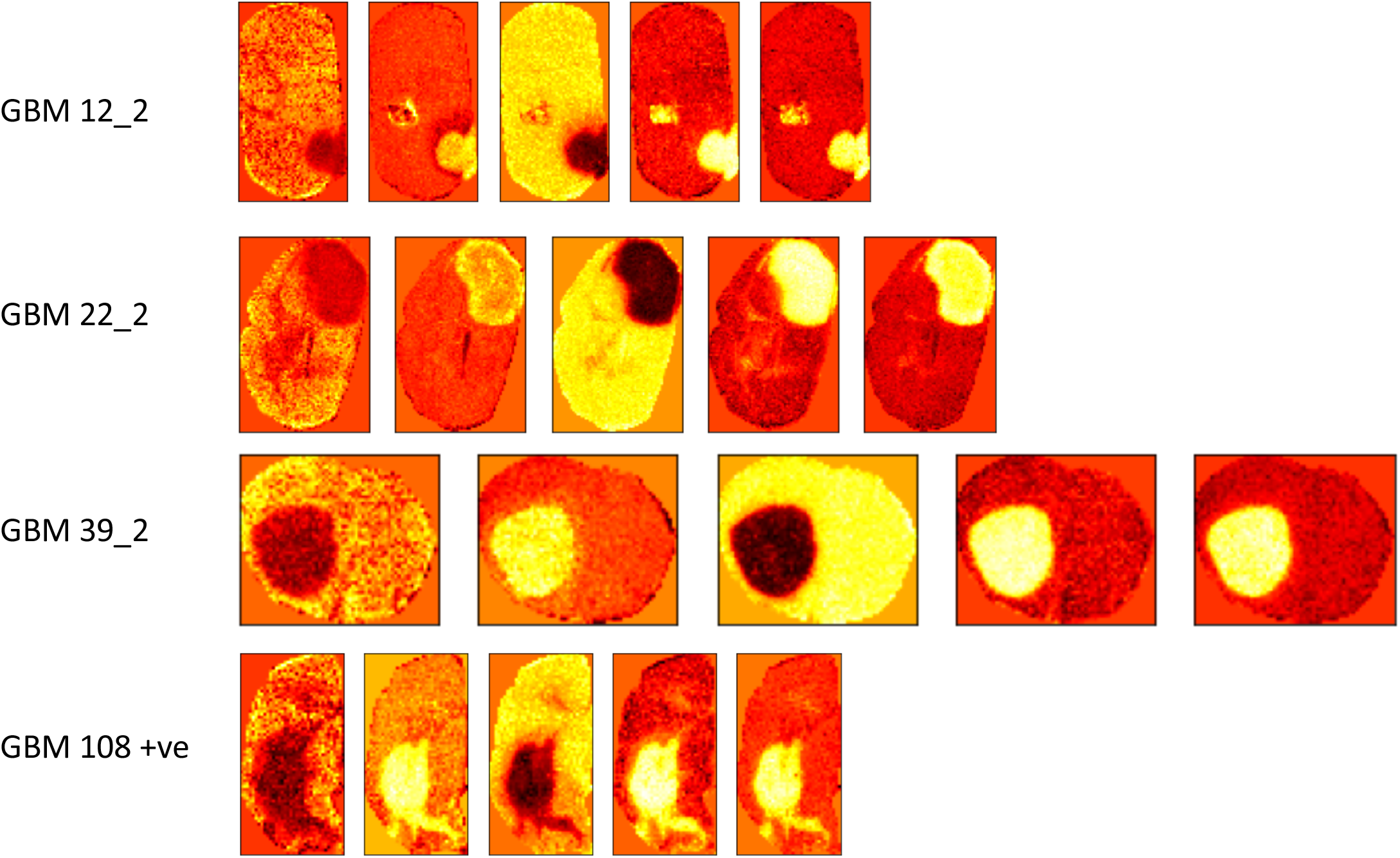
Latent variable of 5-dimensional representation (column) that was captured at the VAE’s code layers for different GBM models (rows) of the MALDI FT-ICR MSI test set. This variable provides a compressed representation of the original MSI data and colormap was arbitrary chosen to reveal structures.

**Table S1.** Evaluation of the classification module using 5-fold cross validation on training/validation set

|  | 1<sup>st</sup> fold | 2<sup>nd</sup> fold | 3<sup>rd</sup> fold | 4<sup>th</sup> fold | 5<sup>th</sup> fold |
| --- | --- | --- | --- | --- | --- |
| Training Accuracy | 1.0 | 1.0 | 1.0 | 1.0 | 1.0 |
| Validation Accuracy | 0.99707 | 1.0 | 1.0 | 1.0 | 1.0 |
| Training Loss | $1.848 \times 10^{-5}$ | $5.474 \times 10^{-6}$ | $3.1 \times 10^{-6}$ | $3.31 \times 10^{-6}$ | $3.345 \times 10^{-6}$ |
| Validation Loss | $8.259 \times 10^{-3}$ | $1.9 \times 10^{-5}$ | $3.488 \times 10^{-6}$ | $3.725 \times 10^{-6}$ | $4.48 \times 10^{-6}$ |

**Table S2.** Classification performance on the training MALDI FT-ICR MSI dataset

|  | class | Spectra # | Precision (%) | Recall (%) | F1-score (%) | Accuracy (%) | Time (min) |
| --- | --- | --- | --- | --- | --- | --- | --- |
| **massNet** | Normal | 10,024 | 100 | 100 | 100 | 100 | 62.2 |
|  | Tumor | 3,644 | 100 | 100 | 100 |  |  |
| **SVM** | Normal | 10,024 | 98 | 99 | 98 | 97.76 | 97.95 |
|  | Tumor | 3,644 | 96 | 96 | 96 |  |  |

**Table S3.** Top 10 correlated m/z peaks with the normal tissue

| *m/z*<br>experimental | correlation | *Tentative*<br>*assignment* | Molecular<br>formula | Adduct | *m/z*<br>calculated | Database | Error<br>(ppm) |
| --- | --- | --- | --- | --- | --- | --- | --- |
| 725.5082 | 0.761263 | PA (36:1) | C39H75O8P | [M+Na]<sup>+</sup> | 725.5091 | Metlin | -1.24 |
| 548.5403 | 0.720929 | Cer (36:1) | C36H71NO3 | [M+H-<br>H<sub>2</sub>O]<sup>+</sup> | 548.5407 | Metlin | -0.73 |
| 552.5069 | 0.719354 |  |  |  |  |  |  |
| 551.5034 | 0.717812 | DG (32:0) | C35H68O5 | [M+H-<br>H<sub>2</sub>O]<sup>+</sup> | 551.5039 | Metlin | -0.91 |
| 553.5097 | 0.713048 |  |  |  |  |  |  |
| 325.76852 | 0.700972 |  |  |  |  |  |  |
| 530.5297 | 0.698111 |  |  |  |  |  |  |
| 699.48376 | 0.696757 | PE-Cer<br>(d34:1) | C36H73N2O6P | [M+K]<sup>+</sup> | 699.4838 | Metlin | -0.06 |
| 580.5383 | 0.696229 |  |  |  |  |  |  |
| 579.53467 | 0.692547 |  |  |  |  |  |  |

## References

Abadi, M. et al. (2016) Tensorflow: A system for large-scale machine learning. 12th {USENIX} Symp. Oper. Syst. Des. Implement. ({OSDI}, 16, 265–283.

Abdelmoula, W.M. et al. (2014) Automatic generic registration of mass spectrometry imaging data to histology using nonlinear stochastic embedding. Anal. Chem., 86.

Abdelmoula, W.M. et al. (2016) Data-driven identification of prognostic tumor subpopulations using spatially mapped t-SNE of Mass spectrometry imaging data. Proc. Natl. Acad. Sci. U. S. A., 113.

Abdelmoula, W.M. et al. (2020) msiPL: Non-linear Manifold and Peak Learning of Mass Spectrometry Imaging Data Using Artificial Neural Networks. bioRxiv Bioinforma., 2020.08.13.250142.

Addie, R.D. et al. (2015) Current State and Future Challenges of Mass Spectrometry Imaging for Clinical Research. Anal. Chem., 87, 6426–6433.

Alexandrov, T. (2012) MALDI imaging mass spectrometry: statistical data analysis and current computational challenges. BMC Bioinformatics.

Alexandrov, T. (2020) Spatial Metabolomics and Imaging Mass Spectrometry in the Age of Artificial Intelligence. Annu. Rev. Biomed. Data Sci., 3, 61–87.

Balluff, B. et al. (2015) De novo discovery of phenotypic intratumour heterogeneity using imaging mass spectrometry. J. Pathol., 235, 3–13.

Basu, S.S. et al. (2019) Rapid MALDI mass spectrometry imaging for surgical pathology. npj Precis. Oncol., 3, 17.

Becht, E. et al. (2019) Dimensionality reduction for visualizing single-cell data using UMAP. Nat. Biotechnol., 37, 38–44.

Behrmann, J. et al. (2018) Deep learning for tumor classification in imaging mass spectrometry. Bioinformatics.

Bowman, A.P. et al. (2020) Ultra-High Mass Resolving Power, Mass Accuracy, and Dynamic Range MALDI Mass Spectrometry Imaging by 21-T FT-ICR MS. Anal. Chem., 92, 3133–3142.

Calligaris, D. et al. (2014) Application of desorption electrospray ionization mass spectrometry imaging in breast cancer margin analysis. Proc. Natl. Acad. Sci., 111, 15184–9.

Calligaris, D. et al. (2015) MALDI mass spectrometry imaging analysis of pituitary adenomas for near-real-time tumor delineation. Proc. Natl. Acad. Sci. U. S. A., 112, 9978–9983.

Caprioli, R.M. (2019) Imaging mass spectrometry: A perspective. J. Biomol. Tech., 30, 7– 11.

Carreira, R.J. et al. (2015) Large-Scale Mass Spectrometry Imaging Investigation of Consequences of Cortical Spreading Depression in a Transgenic Mouse Model of Migraine. J Am Soc Mass Spectrom.

Castellino, S. et al. (2011) MALDI imaging mass spectrometry: Bridging biology and chemistry in drug development. Bioanalysis, 3, 2427–2441.

Chaurand, P. et al. (2004) Integrating Histology and Imaging Mass Spectrometry. Anal. Chem., 76, 1145–1155.

Chollet, F. (2017) Keras (2015). URL http://keras.io.

Coombes, K.R. et al. (2005) Understanding the characteristics of mass spectrometry data through the use of simulation. Cancer Inform., 1, 41–52.

Dewez, F. et al. (2020) MS Imaging-Guided Microproteomics for Spatial Omics on a Single Instrument. Proteomics, 20.

Dexter, A. et al. (2020) Training a neural network to learn other dimensionality reduction removes data size restrictions in bioinformatics and provides a new route to exploring data representations. bioRxiv.

Donnelly, D.P. et al. (2019) Best practices and benchmarks for intact protein analysis for top-down mass spectrometry. Nat. Methods, 16, 587–594.

Drake, R.R. et al. (2017) MALDI Mass Spectrometry Imaging of N-Linked Glycans in Cancer Tissues.

Eberlin, L.S. et al. (2014) Molecular assessment of surgical-resection margins of gastric cancer by mass-spectrometric imaging. Proc. Natl. Acad. Sci. U. S. A., 111, 2436– 2441.

Esteva, A. et al. (2017) Dermatologist-level classification of skin cancer with deep neural networks. Nature, 542, 115–118.

Folk, M. et al. (2011) An overview of the HDF5 technology suite and its applications. In, Proceedings of the EDBT/ICDT 2011 Workshop on Array Databases. ACM., pp. 36– 47.

Guo, D. et al. (2020) Deep multiple instance learning classifies subtissue locations in mass spectrometry images from tissue-level annotations. Bioinformatics, 36, 300–i308.

Heijs, B. et al. (2015) Histology-Guided High-Resolution Matrix-Assisted Laser Desorption Ionization Mass Spectrometry Imaging. Anal. Chem., 87.

Hosny, A. et al. (2018) Artificial intelligence in radiology. Nat. Rev. Cancer, 18, 500–510.

Huizing, L.R.S. et al. (2019) Development and evaluation of matrix application techniques for high throughput mass spectrometry imaging of tissues in the clinic. Clin. Mass Spectrom., 12, 7–15.

Inglese, P. et al. (2017) Variational autoencoders for tissue heterogeneity exploration from (almost) no preprocessed mass spectrometry imaging data. arXiv.

Ioffe, S. and Szegedy, C. (2015) Batch normalization: Accelerating deep network training by reducing internal covariate shift. arXiv Prepr. arXiv1502.03167.

Jones, E.A. et al. (2012) Imaging mass spectrometry statistical analysis. J Proteomics, 75, 4962–4989.

van Kersbergen, J. et al. (2019) Cancer detection in mass spectrometry imaging data by dilated convolutional neural networks. In, Medical Imaging: Digital Pathology (Vol. 10956). International Society for Optics and Photonics., p. 109560I.

Kingma, D.P. and Welling, M. (2013) Auto-encoding variational bayes. arXiv Prepr. 1312.6114.

Lecun, Y. et al. (2015) Deep learning. Nature, 521, 436.

Van Der Maaten, L.J.P. et al. (2009) Dimensionality Reduction: A Comparative Review. J. Mach. Learn. Res., 10, 1–41.

McDonnell Liam A. et al. (2010) Automated imaging MS: Toward high throughput imaging mass spectrometry. J. Proteomics, 73, 1279–1282.

McDonnell, LA et al. (2010) Imaging mass spectrometry data reduction: automated feature identification and extraction. J Am Soc Mass Spectrom, 21, 1969–1978.

McDonnell, L.A. and Heeren, R.M. (2007) Imaging mass spectrometry. Mass Spectrom Rev, 26, 606–643.

McInnes, L. et al. (2018) UMAP: Uniform manifold approximation and projection for dimension reduction. arXiv Prepr. 1802.03426.

Murta, T. et al. (2021) Implications of Peak Selection in the Interpretation of Unsupervised Mass Spectrometry Imaging Data Analyses. Anal. Chem.

Norris, J.L. and Caprioli, R.M. (2013) Imaging mass spectrometry: A new tool for pathology in a molecular age. Proteomics -Clin. Appl., 7, 733–738.

Patterson, N.H. et al. (2018) Next Generation Histology-Directed Imaging Mass Spectrometry Driven by Autofluorescence Microscopy. Anal. Chem., 90, 12404– 12413.

Race, A.M. et al. (2021) Deep Learning-Based Annotation Transfer between Molecular Imaging Modalities: An Automated Workflow for Multimodal Data Integration. Anal. Chem., 93, 3061–3071.

Race, A.M. et al. (2012) Inclusive sharing of mass spectrometry imaging data requires a converter for all. J. Proteomics, 75, 5111–5112.

Randall, E.C. et al. (2018) Integrated mapping of pharmacokinetics and pharmacodynamics in a patient-derived xenograft model of glioblastoma. Nat. Commun., 9.

Randall, E.C. et al. (2020) Localized metabolomic gradients in patient-derived xenograft models of glioblastoma. Cancer Res., 80, 1258–1267.

Ronneberger, O. et al. (2015) U-Net: Convolutional Networks for Biomedical Image Segmentation. Miccai.

Santagata, S. et al. (2014) Intraoperative mass spectrometry mapping of an onco-metabolite to guide brain tumor surgery. Proc. Natl. Acad. Sci., 111, 11121–11126.

Seddiki, K. et al. (2020) Cumulative learning enables convolutional neural network representations for small mass spectrometry data classification. Nat. Commun., 11.

Srivastava, N. et al. (2014) Dropout: A simple way to prevent neural networks from overfitting. J. Mach. Learn. Res., 15, 1929–1958.

Thomas, S.A. et al. (2016) Dimensionality Reduction of Mass Spectrometry Imaging Data using Autoencoders. IEEE Symp. Ser. Comput. Intell., 1–7.

Veselkov, K.A. et al. (2014) Chemo-informatic strategy for imaging mass spectrometry-based hyperspectral profiling of lipid signatures in colorectal cancer. Proc. Natl. Acad. Sci., 111, 1216–1221.

